# A transposable element insertion is the switch between alternative life history strategies

**DOI:** 10.1101/424879

**Authors:** Alyssa Woronik, Kalle Tunström, Michael W. Perry, Ramprasad Neethiraj, Constanti Stefanescu, Maria de la Paz Celorio-Mancera, Oskar Brattström, Jason Hill, Philipp Lehmann, Reijo Käkelä, Christopher W. Wheat

## Abstract

Tradeoffs affect resource allocation during development and result in fitness consequences that drive the evolution of life history strategies. Yet despite their importance, we know little about the mechanisms underlying life history tradeoffs in wild populations. Many species of *Colias* butterflies exhibit an alternative life history strategy (ALHS) where females divert resources from wing pigment synthesis to reproductive and somatic development. Due to this reallocation, a wing color polymorphism is associated with the ALHS: individuals have either yellow/orange or white wings. Here we map the genetic basis of the ALHS switch in *Colias crocea* to a transposable element insertion downstream of the *Colias* homolog of *BarH-1*, a homeobox transcription factor. Using CRISPR/Cas9 gene editing, antibody staining, and electron microscopy we find morph-specific specific expression of *BarH-1* suppresses the formation of pigment granules in wing scales. Lipid and transcriptome analyses reveal physiological differences associated with the ALHS. These findings characterize a novel mechanism for a female-limited ALHS and show that the switch arises via recruitment of a transcription factor previously known for its function in cell fate determination in pigment cells of the retina.

---

A life-history strategy is a complex pattern of co-evolved life history traits (e.g. number of offspring, size of offspring, and lifespan^1^), that is fundamentally shaped by tradeoffs that arise because all fitness components cannot simultaneously be maximized. Therefore, finite resources are competitively allocated to one life history trait versus another within a single individual, and selection acts on these allocation patterns to optimize fitness^2^. Evolutionary theory predicts that positive selection will remove variation from natural populations, as genotypes with the highest fitness go to fixation^3^. However, across diverse taxa alternative life history strategies (ALHSs) are maintained within populations at intermediate frequencies due to balancing selection^4^. Life history theory was developed using methods such as quantitative genetics, artificial selection, demography, and modeling to gain significant insights into the causes and consequences of genetic and environmental variation on life history traits. Yet despite these advances, a key challenge that remains is to identify the proximate mechanisms underlying tradeoffs, especially for ecologically relevant tradeoffs that occur in natural populations^5^. Here, we identify the mechanism underlying one such ALHS in the butterfly *Colias crocea* (Pieridae, Lepidoptera) (Geoffroy, 1785).

*Colias* butterflies (the “clouded sulphurs”) are common throughout the Holarctic and can be found on every continent except Australia and Antarctica^6^. In approximately a third of the nearly 90 species within the genus, females exhibit two alternative wing-color morphs: yellow or orange (depending on the species) and white^6, 7^ (Fig. 1A). The wing color polymorphism arises because during pupation the white morph, also known as Alba, reallocates larval derived resources from the synthesis of energetically expensive colored pigments to reproductive and somatic development^8^. This tradeoff has been well characterized in *Colias crocea*, the Old World species that we focus upon in this work, via radio-labelled metabolite tracking in pupae^9^ as well as in the New World species *Colias eurytheme^8^* (Pieridae, Lepidoptera) (Boisduval, 1852) using ultraviolet spectrophotometry. As a result of the resource reallocation, Alba females have faster pupal development, a larger fat body, and significantly more mature eggs at eclosion compared to orange females^10^. However, despite these developmental advantages and the dominance of the Alba allele, the polymorphism is maintained by several abiotic and biotic factors ^10–14^. For example, males preferentially mate with orange females, as wing color is an important cue for mate recognition^10,12,13^. This mating bias likely has significant fitness costs for Alba females because males transfer essential nutrients during mating, and multiply mated females have more offspring over their lifetime^15, 16^. The mating bias against Alba females is strongest in populations that frequently co-occur with other white Pierid butterfly species due to interference competition^13^. Also, Alba’s development rate advantage is temperature dependent, with Alba females having faster development in cold temperatures^10^. Field studies confirm Alba frequency and fitness increases in species that inhabit cold and nutrient poor habitats, where the occurrence of other white Pierid butterflies is low. While in warm environments with nutrient rich host plants and a high co-occurrence of other white species, orange females exhibit increased fitness and frequency^12–14^. Previous work has also suggested Alba females have a higher sensitivity to viral infections^9^. In all *Colias* species where it has been investigated (n=6), the switch between the Alba or the orange strategy is controlled by a single, autosomal locus^6^. This fact, along with ancestral state reconstruction^7^, has led to the assumption that the Alba locus is conserved within the genus *Colias*, and potentially across the subfamily Coliadinae. Yet, despite over a century of research on various aspects of Alba biology the mechanism underlying this polymorphism remained unknown.

**Fig. 1.**
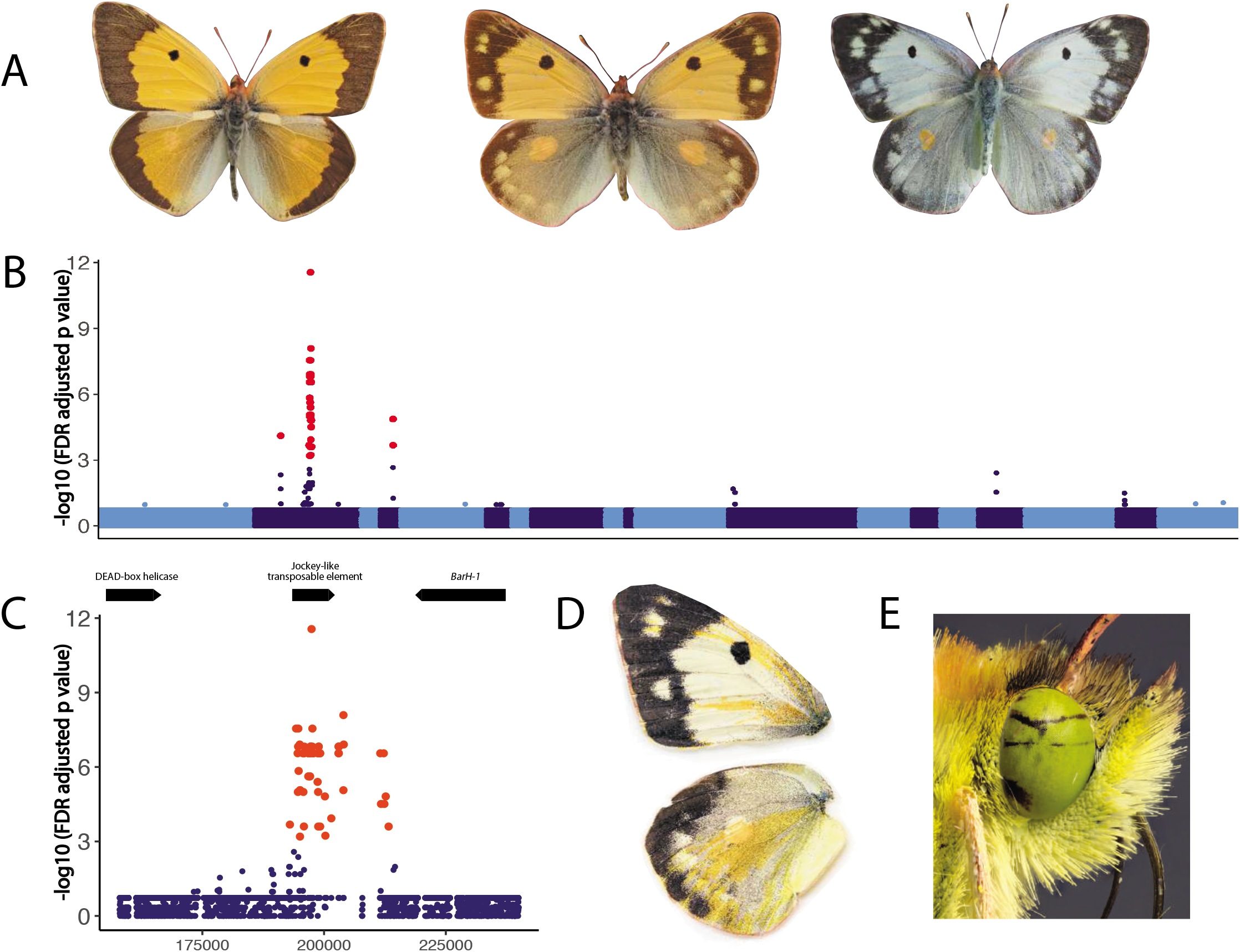
Color variation in *Colias crocea* and the genetic mechanism of Alba. (A) *Colias crocea* male, orange female, and Alba female (left to right). (B) SNPs significantly associated with the Alba phenotype (red) within the ~3.7 Mbp Alba locus identified via 3 rounds of bulk segregant analysis. Contigs in this region shown as alternating dark and light blue. (C) The location of Alba associated SNPs (red) on the ~430 kb outlier contig identified in the GWAS. Gene models for the DEAD-box helicase, the Jockey-like transposable element, and *BarH-1* shown at the top of the panel. (D) Wings of a female with an Alba genotype following CRISPR/Cas9 mosaic knockout of *BarH-1*, wild type regions are white, knockout regions are orange. Orange color is seen on the dorsal forewing (top) and hindwing (bottom). (D) *BarH-1* mosaic knockout also leads to black regions in the eyes, wild type regions are green. !

Using a *de novo* reference genome for *C. crocea* that we generated via Illumina and PacBio sequencing, and three rounds of bulk segregant analyses (BSA) using whole genome sequencing from a female and two male informative crosses for Alba, we mapped the Alba locus to a ~3.7 Mbp region (Supplementary Fig. 1, & Supplementary Information). Then, with whole genome re-sequencing data from 15 Alba and 15 orange females from diverse population backgrounds, a SNP association study fine mapped the Alba locus to a ~430 kb contig that fell within the ~3.7 Mbp locus identified using the BSA crosses (Fig. 1B and Supplementary Information). The majority of SNPs significantly associated with Alba (n=70 of 72) were within or flanking a *Jockey-like* transposable element (TE) (Fig. 1C). We determined that the TE insertion was unique to the Alba morph in *C. crocea* by assembling orange and Alba haplotypes for this region, then quantifying differences in read depth between morphs within and flanking the insertion (Supplementary Information and Supplementary Figs. 2, 3, & 4). We then used PCR to validate the presence or absence, respectively, of the insertion in 25 Alba and 57 orange wild-caught females (Supplementary Fig. 7). We also found no evidence of a TE insertion in the homologous region of other butterfly genomes *(Danaus plexippus* & *Heliconius melpomene)* (Supplementary Fig. 2).

The Alba-specific insertion was located ~30 kb upstream of a gene encoding a DEAD-box helicase, and ~6kb downstream of the *Colias* homolog of *BarH-1*, a homeobox transcription factor (Fig. 1C). *BarH-1* was an intriguing find as it affects color via pigment granule development within eyes of *Drosophila melanogaster^17^*. To investigate BarH-1 expression in developing *C. crocea* wings, we used *in situ* hybridization of BarH-1 on wings from two day old pupae of orange and Alba females. We found the BarH-1 protein is expressed in scale building cells within the white wing regions in Alba females (Fig. 2B). We did not observe BarH-1 in scale building cells from orange areas of the wing in orange females (Fig, 2C). Interestingly however, we found BarH-1 is expressed in scale building cells within black regions for both morphs (Fig, 2A&D). To validate the functional role of *BarH-1* in the Alba phenotype, we generated CRISPR/Cas9-mediated deletions within exons 1 and 2 using a mosaic knockout (KO) approach (Supplementary Information). *BarH-1* KO gave rise to a white/orange color mosaic on the dorsal side of the wings in females with an Alba genotype (i.e. TE insertion +) (Fig. 1D), while KO males and orange females displayed no white/orange mosaic on the wing. These results indicate *BarH-1* expression suppresses orange coloration in the wings. We also observed black and green mosaic coloring of eyes in KO males and females of both morphs, where green eyes are the wild type color (Fig. 1E). These results indicate *BarH-1* also plays a role in *Colias* eye development.

**Fig. 2.**
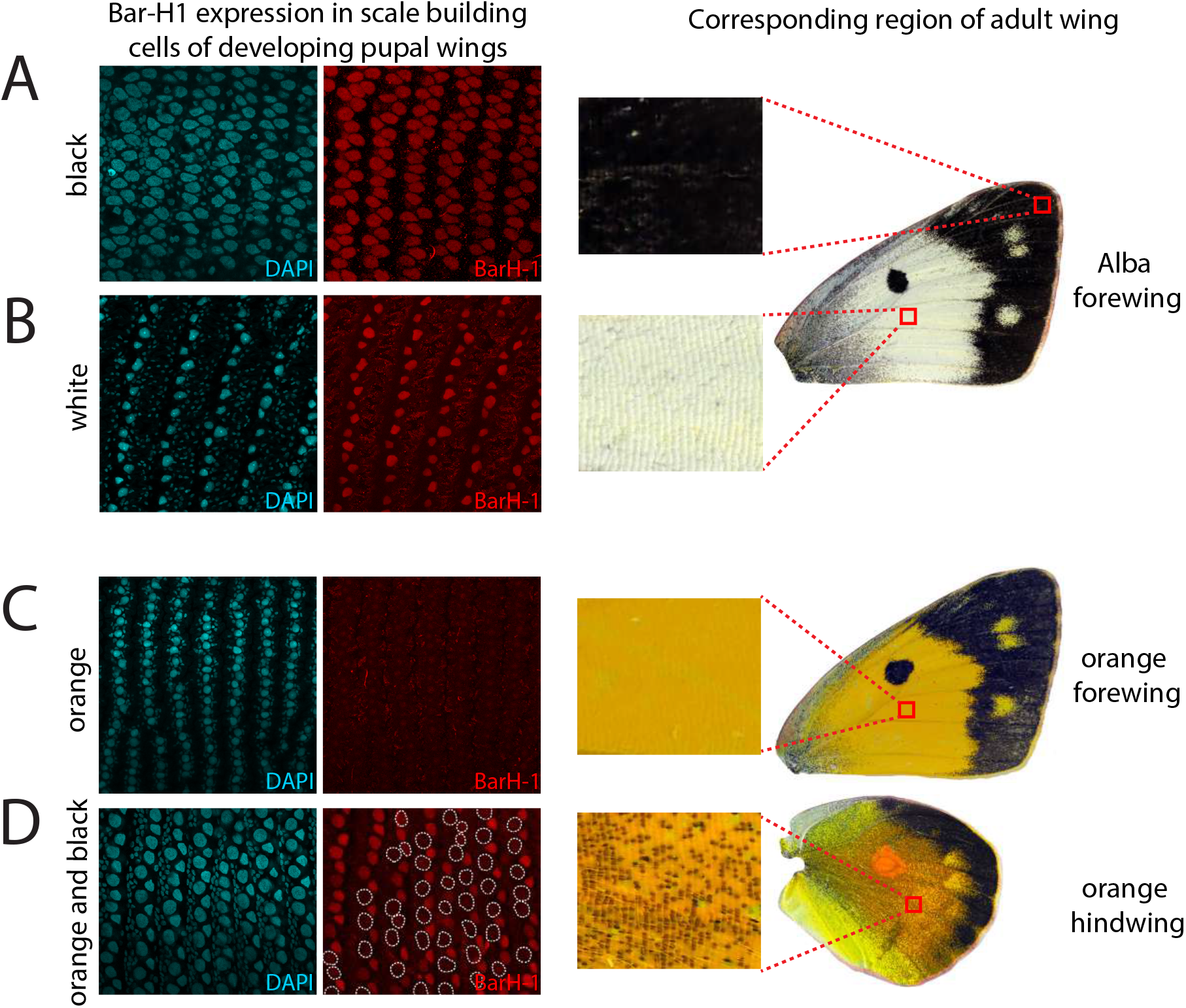
BarH-1 is expressed in white but not orange regions of the wing in *C. crocea* females. DAPI (nuclei, left, blue) and BarH-1 antibody (right, red) staining of pupal wings. Large nuclei are in scale building cells, small nuclei are in epithelial cells. The right part of the panel shows the approximate location of the stained area and the scales in this region in an adult wing. Scale bars are 2μm. (A) Staining of the forewing of an Alba female in an area at the black margin (top) and a white area (bottom). BarH-1 is expressed in melanic as well as white Alba scale building cells. (B) Antibody staining of the forewing of an orange female. BarH-1 is not expressed in these scale building cells. (C) Antibody staining of the hindwing of an orange female. BarH-1 is heterogeneously expressed in the scale building cells within this region. This staining pattern presumably corresponds to the variation in scale color, with melanic scale building cells expressing BarH-1 but orange lacking expression.

We next investigated how the Alba color change manifests within wings. Butterfly wing color can arise either due to the absorption of light by pigments deposited within the scales, or by the scattering of light via regularly arranged nanostructures in the scales^18^. *Colias* butterflies have pteridine pigments. These pigments are synthesized within the wings and previous work using ultraviolet spectrophotometry in *C. eurytheme* found Alba females exhibit dramatic reductions in colored pteridine pigments compared to orange^8, 9^. In insects, pteridines are synthsized in pigment granules and pigment granules are concentrated within wing scales of Pierid butterflies^19, 20^. However, whether morphs differed in wing scale morphology was unknown. To investigate wing morphology, we used scanning electron microscopy and found white scales from Alba individuals exhibited a dramatic and significant reduction in pigment granules, compared to orange scales (t_5.97_ = 2.93, p = 0.03) (Fig 3 A&B). These results indicate the color change to white is caused by reduced pigment granule formation. Congruent with this interpretation, CRISPR KO Alba individuals exhibited significantly less pigment granules in scales from the white wild-type region compared to scales in orange *BarH-1* KO regions (t_5.45_ = 10.78, p < 0.001) (Fig. 3C). To further test whether reduction in pigment granule amount alone was sufficient for the orange to white color change, we chemically removed the pigment granules from the wing of an orange *C. crocea* female. This resulted in formerly orange regions turning white (Fig. 3D). Wings likely appear white after granule removal due to the scattering of light from the remaining non-lamellar nanosctructures^21^. These results demonstrate that *BarH-1* suppresses pigment granule formation in wing scales, resulting in the white color of Alba females in *C. crocea*. Thus, we propose the resource tradeoff between color and development arises due to a classic Y reallocation model, wherein limited resources are competatively allocated and increased investement in one trait results in a decreased investment to another^22^. Within the energetically closed system of a developing pupa, reduced pigment granule formation would likely result in reduced pigment synthesis, which would in turn leave more resources free to be used for other developmental processes. Finally, we also observed scale building cells in black regions of both morphs express BarH-1 and also lack pigment granules (Fig 2 A&D and Fig 3 A&B), but these scales appear black due to melanin deposition within the scale^18^. These results suggest BarH-1 may also repress pigment granule formation within black scales.

**Fig. 3.**
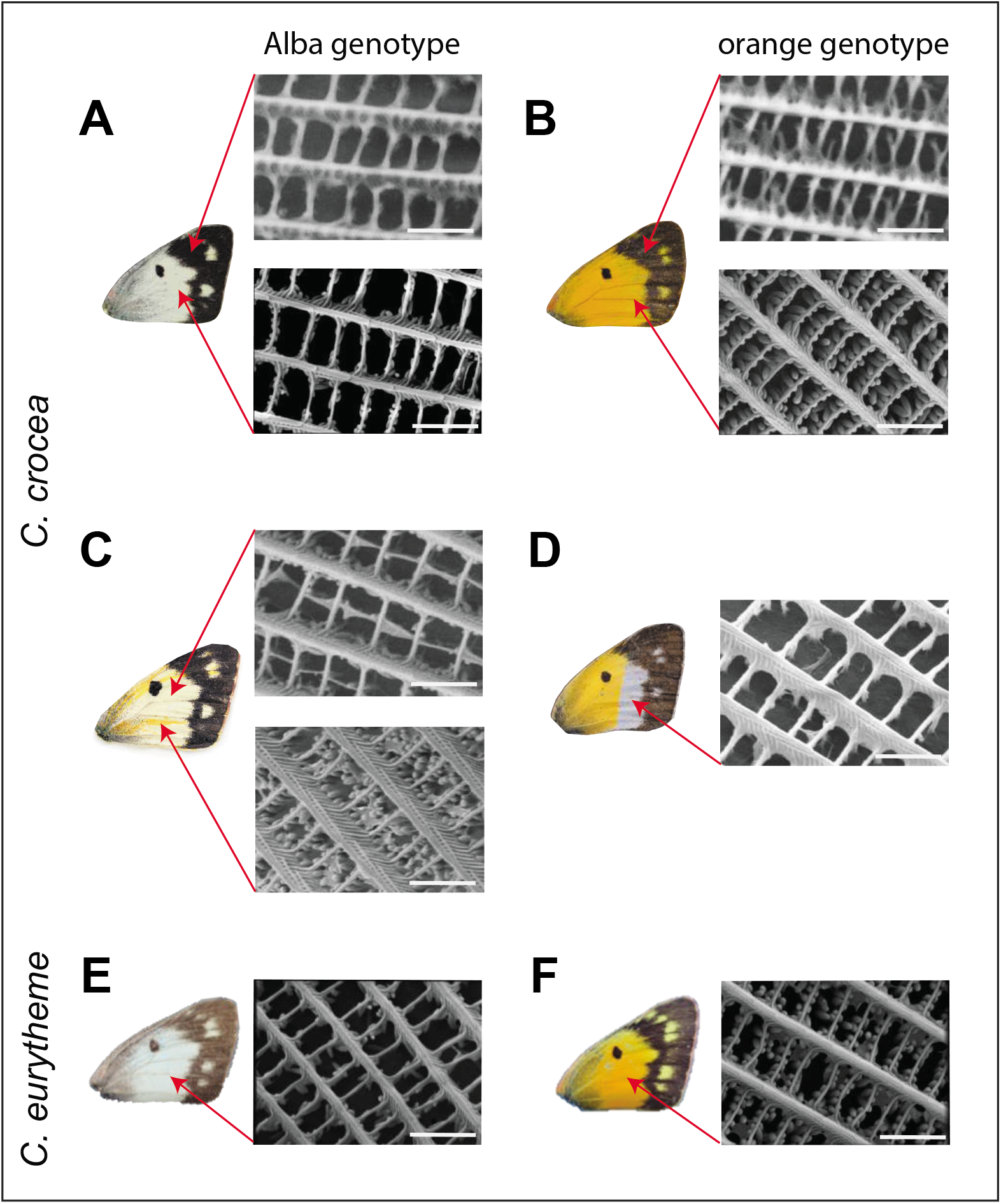
*Colias* forewings and scanning electron microscope (SEM) images of their wing scale nanostructures. (A) *C. crocea* Alba female wing and wing scale structure. The top panel shows the SEM image of a black scale; pigment granules are absent. The bottom panel shows a white scale, exhibiting near absence of pigment granules. (B) Wing and wing scale structures of a wild type orange *C. crocea* female. The top panel shows a black scale, pigment granules are absent. The bottom panel shows an orange scale with abundant pigment granules. (C) Wing and wing scales of a genetically Alba female exhibiting CRISPR/Cas9 mosaic knockout of *BarH-1*. The top panel shows the wild-type white scale, where pigment granules are mostly absent. The bottom panel shows a scale in an orange *BarH-1* KO region. It exhibits significantly more pigment granules than the white scales. (D) Wing and wing scales of an orange *C. crocea* female where pigment granules have been chemically removed from the distal half of the wing. The SEM image shows a scale from the white region with pigment granules completely missing. The white color of this wing section presumably results from light reflection off the remaining scale nanostructures. (E) Wing and wing scale structure of a *C. eurytheme* Alba female. Wing scales lack pigment granules, similar to the phenotype observed in *C. crocea*. (F) Wing and wing scale structures of a *C. eurytheme* orange female. Orange scales show abundant pigment granules, again consistent with the orange phenotype observed in *C. crocea*.

The Alba mechanism is assumed to be conserved across *Colias*. Therefore, we wished to test whether Alba females from the New World species *Colias eurytheme* also exhibited significantly less pigment granules than orange females. Indeed, we found orange *C. eurytheme* scales exhibited abundant pigment granules while Alba scales almost entirely lacked granules (Fig. 3 E&F). These results demonstrate white wing color arises via the same morphological mechanism within *Colias* and corroborate previous assumptions that Alba is conserved across the genus. To further validate that other aspects of the Alba/orange alternative life history strategy are conserved across the genus we tested whether one of the physiological tradeoffs of Alba reported for New World species was also seen in *C. crocea*. In *C. eurytheme*, Alba females have larger fat bodies than orange females and the strength of the Alba advantage increased in cold temperatures^10^. To compare abdominal lipid stores between morphs in *C. crocea*, we conducted high performance thin layer chromatography on two day old adult females reared under two temperature treatments (Hot: 27°C vs. Cold: 15°C during pupal development). Adults were not allowed to feed before samples were taken, therefore these measurements reflect larval stores, where the putative energetic tradeoff should be more clearly visible. We found Alba females had larger abdominal lipid stores than orange in both temperature treatments, though the difference was only significant in the cold treatment (cold: n=32, t_29.12_ = 3.42, P = 0.002, hot: n=25, t_22.71_ = 0.67, P = 0.51) (Fig. 4A). These results are consistent with previous reports from New World *Colias* species and indicate that the morph-specific tradeoff associated with the color change is also conserved across the genus.

**Fig. 4.**
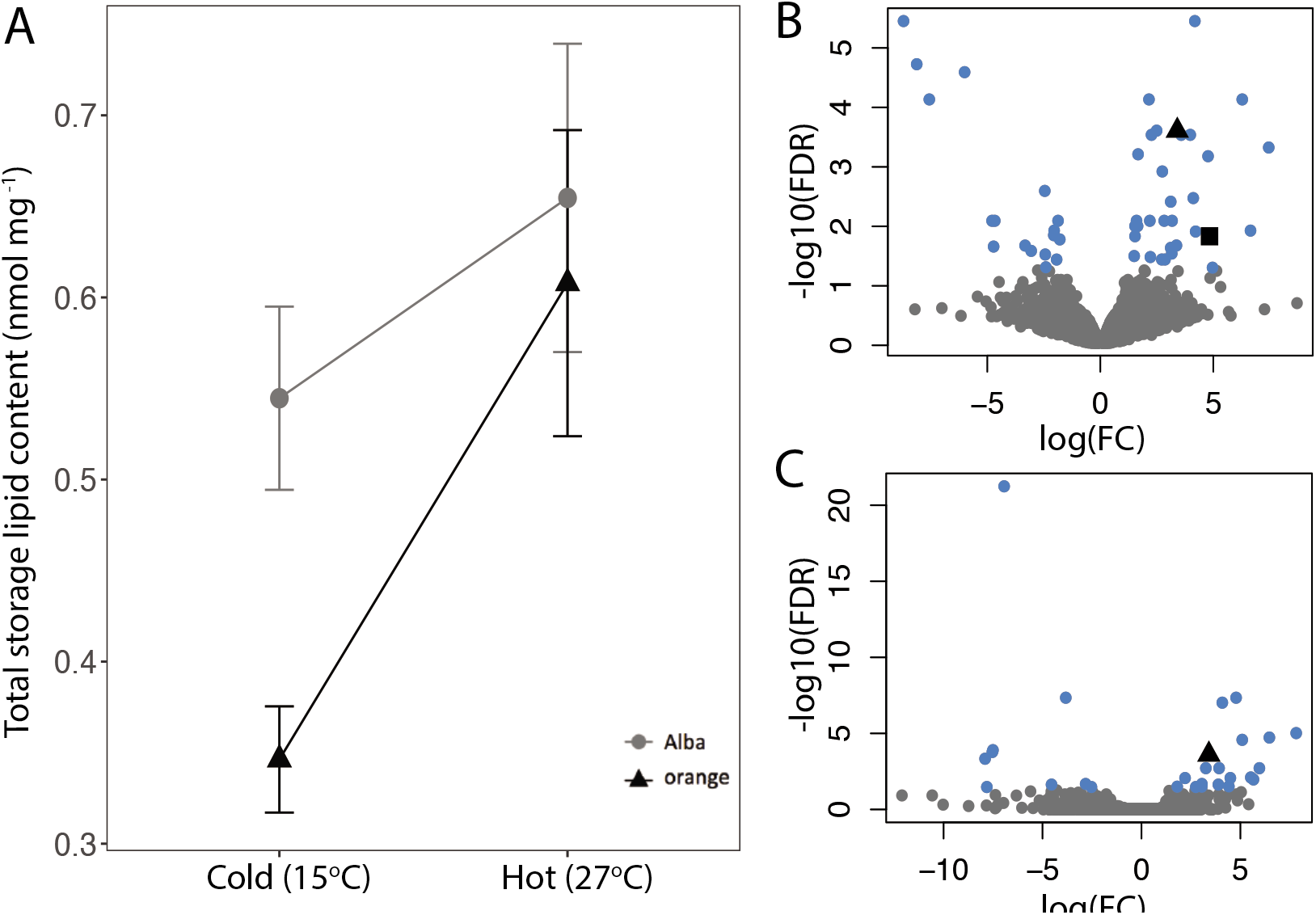
Physiological differences between female morphs of *C. crocea*. A) The mass corrected total neutral lipid content for female morphs in two temperature treatments. Alba females, on average, have larger neutral lipid stores than orange females. However there is an interaction between morph and temperature as the difference is only significant in the cold treatment. Error bars are the standard error (cold: n=32, t_29.12_ = 3.42, P = 0.002, hot: n=25, t_22.71_ = 0.67, P = 0.51). B) Volcano plot to visualize gene expression differences between female morphs in pupal abdominal tissue. Each point is a gene. Genes not significantly differentially expressed between morphs are grey, while differentially expressed genes are blue. The black square is the triacylglycerol lipase and the black triangle is *RIM*. The X-axis is the log of the fold change (FC), positive log(FC) indicates the gene is upregulated in Alba individuals. C) Volcano plots to visualize gene expression differences between female morphs in pupal wing tissue. Color coding, shapes, and axes are the same as above.

We then investigated the transcriptome of pupal abdomen and wing tissue at the time of pteridine synthesis to identify genes that exhibited differential expression between morphs and therefore may play a role in the morph-specific differences in physiology that arise due to the resource tradeoff (Fig. 4 B&C, Supplementary Information, Supplementary Tables 5,6,&7). In *C. eurytheme* Alba females emerge from the pupa with significantly more mature eggs than orange females^10^ and we find evidence that suggests similar dynamics are occurring in *C. crocea*. A gene set enrichment analysis (GSEA) revealed that ‘embryo development ending in birth or egg hatching’ (GO:0009792, p = 0.00072), ‘proteasome-mediated ubiquitin-dependent protein catabolic process’ (GO:0043161, p = 0.00073), and ‘proteolysis’ (GO:0006508, p = 0.00101) were within the top 5 terms enriched and upregulated within Alba abdomens (Supplementary Table 6). Additionally our differential expression analysis identified a that gene which encodes a triacylglycerol lipase was significantly upregulated within Alba abdomen tissue (log fold change [log FC] of 4.8) (Fig. 4B and Supplementary Table 5). Triacylglycerol composes more than 90% of the lipids stored in the fat body and during times of energy demand triacylglycerol lipases mobilize these stores^23^. For example, during embryogenesis there is a massive shift in lipid distribution from the fat body to ovaries as lipids comprise 30-40% of the dry weight of insect embryos^23^. Taken together these results suggest that, similar to *C. eurytheme*, Alba females of *C. crocea* may be benefitting from increased embryogenesis compared to orange females. We also observe an enrichment of ‘defense response to Gram-positive bacterium’ (GO:0050830, 0.00027) for genes upregulated within Alba abdomens. Interestingly, previous work has suggested that Alba females may have enhanced sensitivity to viral infection^9^. Further investigation of potential morph-specific tradeoffs between wing color and immunity is of interest.

For genes downregulated in Alba abdomens the GSEA revealed significant enrichment of ‘regulation of nucleoside metabolic process’ (GO:0009118, p value < 0.0001) and ‘regulation of purine nucleotide catabolic process’ (GO:0033121, p value < 0.0001) (Supplementary Table 6). *Colias* wings use purine precursors to synthesize pteridines^24^. Downregulation of these GO terms in Alba abdomens suggests that the decreased pteridine synthesis observed in Alba females^8^, which likely arises due to the decrease in pigment granules in the wings, leads to a decrease in purine precursors being shunted from the abdomen to the wings. Additionally, consistent with previous reports of GTP reallocation from wings to other areas of development in Alba females^8^ we also observed significant enrichment of ‘positive regulation of GTPase activity’ (GO:0043547, p value < 0.0001). Additionally, *RIM*, a Rab GTPase effector^25^, was one of the most highly differentially expressed (DE) genes in both tissues (logFC increase in Alba of 3.4 in the abdomen and 5.1 in the wings) (Fig. 4 B&C and Supplementary Table 5). RIM acts as a molecular switch by converting guanosine diphosphate to guanosine triphosphate (GTP), thereby activating its associated Rab GTPase, which is in turn involved in synaptic vesicle exocytosis and secretory pathways^26^.

Within wings, *BarH-1* was not differentially expressed between morphs at this stage, indicating that morph specific expression differences are temporal. However, we did observe genes downregulated in Alba wings were significantly enriched for ‘xanthine dehydrogenase activity’ (p = 0.02, GO:0004854) (Supplementary Table 7). Xanthine dehydrogenase is the enzyme that catalyzes the xanthopterin to leucopterin conversion during pteridine synthesis in *Colias* butterflies^8^. These results are consistent with previous studies in *C. eurytheme* that reported the level of xanthopterin in Alba wings was 7-8 fold less than in orange^8^. Additionally we observed enrichment of ‘MAP kinase activity’ (GO:0004709, p = 0.00109) in genes downregulated within Alba wings. In *Drosophila*, BarH-1 represses Decapentaplegic a morphogen that is homolog to TGFß^27^. TGFß can activate signalling cascades, including MAP kinase pathways^28^. Previous work in *Drosophila* has also suggested an interaction between Bar homeobox genes and Ras/MAP kinase signalling during eye development^29^. Future functional studies of candidate genes are needed to better understand their mechanistic roles in wing development and the tradeoffs associated with the ALHS.

Here we identified the proximate mechanism underlying a female-limited ALHS in a natural population. Historically, the field of life history research has treated mechanistic details as a black box^5^, though recently several genetic mechanisms underlying ecologically relevant ALHSs have been identified, e.g. in the wall lizard^30^, ruff^31, 32^, white throated sparrow^33^, and fire ant^34^. The majority of these studies found that supergenes, large loci that maintain many genes in tight linkage due to structural variation, gave rise to the alternative morphs^31–35^. Such findings established that structural variation facilitates the evolution of complex traits. However these genomic architectures make determining the specific contributions of individual genes to ALHSs difficult, though there have been significant advances made in the white throated sparrow^33^. Interestingly, our results, and recent work in the wall lizard^30^, found that ALHSs arose due to changes in the regulatory region of either a single or two genes, respectively. This raises the question of how these regions give rise to the other fitness-related traits associated with the ALHS. We propose the Alba-associated physiological and developmental traits arise due to a classic Y reallocation model, where reduced pigment granule formation results in reduced pigment synthesis, which in turn leaves more resources free to be used for other developmental processes. Previous literature studying the Alba phenotype had been unable to determine whether allocation of resources from the fatbody, or pigment biosynthesis within the wing scales, was the basis of Alba. Here, the mosaicism of the BarH-1 KO documents the cell level autonomy of the Alba polymorphism, as abdomen level provisioning would affect all wing scales equally. Nevertheless, there may be other pleiotropic effects of the reallocation, such as the sensitivity of the Alba to viral disease, or of the TE insertion itself as it may affect BarH-1 expression in tissues other than the wing.

Previous work has shown that *BarH-1* plays a role in the morphogenesis of neurons, leg segments, and eyes in *Drosophila*^36^. Specifically, *BarH-1* expression is required for the formation of pigment granules and the deposition of red pteridine pigments in the *Drosophila eye^17^*. We find that *BarH-1* also plays a role in eye and wing color in *Colias* butterflies. However, as *BarH-1* expression represses the formation of pigment granules within *Colias* wings, we find it has a reversed function in *Drosophila* and *Colias*. This may be one of several examples where either whole or a part of a gene regulatory network that regulates eye development has been co-opted to give rise to a novel trait in the insect wing^37^. If so, future work could investigate what aspects of the network have been co-opted and how this lead to BarH-1’s contrasting roles in morphogenesis.

Additionally, recent work in the field of butterfly wing evolutionary-development has found that several genes are repeatedly involved in wing color variation across distantly related species. Such genes form a patterning “toolkit” (e.g. *optix*^38^, *WntA*^39^, and *cortex*^40^). *BarH-1* might serve as another toolkit gene for patterning wing color in butterflies beyond *Colias* as we found BarH-1 expression in scale building and socket cells of developing wings in *Vanessa cardui* (Nymphalidae, Lepidoptera) (Linnaeus, 1758) pupae (Supplementary Fig. 10). However, the functional role of BarH-1 in *V. cardui* wings remains to be determined. BarH-1 may have a novel function within *V. cardui* wings. Alternatively, the function of BarH-1 as a repressor of pigment granule formation could be conserved, as *V. cardui* scales do not have pigment pigment granules^41^

Under the latter assumption, we would expect that BarH-1 is not expressed in closely related Pierinae species that, despite appearing white, exhibit abundant pigment granules that are primarily filled with the UV-absorbing pteridine called leucopterin^42^. Future work investigating the evolutionary history of BarH-1’s co-option to the wing and function in other species could shed light on how complex traits such as ALHSs evolve.

## Author Contributions

AW conducted butterfly rearings and lab work, analysed the data, and wrote the manuscript with CWW and input from the coauthors. AW, MWP, KT, and CWW conducted the CRISPR/Cas9 knockout experiment. AW and KT conducted the electron microscopy. MWP conducted antibody staining. RN and JH assisted with bioinformatics. PL and RK conducted HPTLC and AW and PL analyzed the data. AW, CS, CWW and OB conducted fieldwork. MC conducted lab work. CWW supervised the work at all stages.

## Acknowledgements

We would like to thank Lovisa Wennerström, Elishia Harji, Jofre Carnicer, and Christina Hansen Wheat for help with fieldwork. We thank Marianne Ahlbom for assistance with the SEM. Finally we would like to thank Karin Kiontke, Christen Bossu, Naomi Keehnen, and Peter Pruisscher for helpful comments on the manuscript. We thank the Department of Zoology at Stockholm University, the Swedish Research Council 2012–3715, the Academy of Finland 131155, the Knut and Alice Wallenberg Foundation 2012.0058 and the Erik Philip-Sörensens foundation for funding.

## Methods

For detailed methods, including all bioinformatic commands, please see the supplementary information.

### Data availability

SRA reference numbers for the genome and sequencing data will be included upon acceptance.

### Genome assembly

An orange female and male carrying Alba (offspring from wild caught butterflies, Catalonia, Spain) were mated in the lab. DNA from an Alba female offspring of this cross was extracted. Quality and quantity were assessed using a Nanodrop 8000 spectrophotometer (Thermo Scientific) and a Qubit 2.0 fluorometer (dsDNA BR, Invitrogen). A 180 insert size paired end library (101bp reads) was prepared (TruSeq PCR free) and sequenced on an Illumina Hiseq 4000 at the Beijing Genomics Institute (Shenzhen, China). A Nextera mate-pair library with a 3 kb insert size was prepared and sequenced on an Illumina HiSeq 2500 (125bp reads) at the Science for Life Laboratory (Stockholm, Sweden). Raw data was cleaned and high quality reads were used as input for the AllPaths-LG (v. 50960)^43^ assembly pipeline. High molecular weight DNA was extracted from two more Alba females from the above mentioned cross (i.e full siblings). Equal amounts of DNA from each individual were pooled sent to the Science for Life Laboratory (Stockholm, Sweden) for PacBio sequencing on 24 SMRT cells (~17GB of data was produced). A Falcon (v. 0.4.2)^44^ assembly was generated by the Science for Life Laboratory. We then used Metassembler (v. 1.5)^45^ to merge our AllPathsLG and Falcon assemblies, using the AllPathsLG assembly as the primary assembly.

### Bulk segregant analyses (BSA)

The female informative cross data and mapping protocol described in Woronik and Wheat, 2017^46^ was applied to the high quality reference genome to identify the contigs that made up the Alba chromosome. Male Informative Cross (MIC) I: DNA was extracted from a wild caught orange mother (Catalonia Spain) and 26 of her Alba and 24 of her orange female offspring. DNA quality and quantity of each individual was assessed via a Nanodrop 8000 spectrophotometer (Thermo Scientific, MA, USA) and a Qubit 2.0 Fluorometer (dsDNA BR; Invitrogen, Carlsbad, CA, USA) before pooling equal amounts of high-quality DNA from Alba and orange offspring into two pools, respectively. The library preparation (TruSeq PCR-free) and Illumina sequencing (101 bp PE HiSeq2500), was performed at the Beijing Genomics Institute (Shenzhen, China). Raw reads were cleaned and then mapped to the reference genome using NextGenMap v0.4.10 (-i 0.09)^47^. SAMTOOLS v1.2^48^ was used to filter (view -f 3 -q 20), sort and index the bam files and generate mpileup files for the two pools and the orange mother. Popoolation2^49^ were used to calculate the allele frequency difference between Alba and orange pools. SNP sites were filtered in R^50^, for a read depth ≥ 30 and ≤ 300, a bi-allelic state, and a minimum minor allele frequency of 3. The orange mother mpileup was similarly analyzed using Popoolation^51^ (read depth ≥ 5 and ≤30); but the major and minor allele frequencies were calculated in R^50^ by dividing the major and minor allele count by the read depth at each site respectively. A SNP site was considered a MIC I Alba SNP when it met the following expectations: 1) homozygous in the orange mother, 2) homozygous in the orange pool, 3) the allele frequency difference in the Alba pool compared to the orange was 0.45-0.55. MIC II: A male carrying Alba mated an orange female in the lab at Stockholm University. DNA was prepared as described above for 26 Alba and 28 orange female offspring resulting in two DNA pools. Library preparation (TruSeq PCR-free) and Illumina sequencing (150 bp paired-end reads with 350bp insert, HiSeqX), was performed at Science for Life Laboratory (Stockholm, Sweden). The same mapping and SNP calling pipeline used on the MIC I was applied. A site was considered an Alba SNP if 1) it was homozygous in the orange pool and 2) the allele frequency difference in the Alba pool compared to the orange was 0.45-0.55. A contig was considered Alba associated if it had ≥ 3 Alba SNPs in all crosses. Nineteen Alba associated contig were identified. They total ~3.7Mbp and are considered the Alba BSA locus.

### Genome wide association study

DNA for genome re-sequencing was extracted from 15 Alba and 15 orange females from diverse population backgrounds (Catalonia, Spain and Capri, Italy). High quality DNA was prepared using Illumina TruSeq and sequenced at the Science for Life Laboratory (Stockholm, Sweden) (150 bp paired-end reads HiSeqX). Cleaned reads were mapped to the annotated reference genome using NextGenMap v0.4.10 (-i 0.6 -X 2000)^47^. Bam files were filtered and sorted using SAMTOOLS v1.2 (view -f 3 -q 20) ^48^. A VCF file was generated using SAMTOOLS v1.2 (-t DP -t SP -Q 15)^48^ and bcftools v. 1.2 (-Ov -m) ^48^. VCFtools (v0.1.13)^52^ was used to call SNP sites with no more than 50% missing data, an average read depth between 15-50 across individuals, and a minimum SNP quality of 30. An association analysis was performed with PLINK (v1.07)^53^ and a Benjamini & Hochberg step-up FDR control was applied. SNPs with FDR <0.05 were considered Alba SNPs. We conducted this analysis both genome wide and only within the BSA locus. Both analyses fine mapped the Alba locus to the same genomic region.

### Antibody Generation and Staining

A Rabbit-anti-Bar antibody was generated against the full length sequence of the *Vanessa cardui* Bar homolog. Protein was generated by GenScript (Piscataway, NJ) and purified to >80% purity. DNA sequences to produce this protein were codon-optimized for bacterial expression and made via gene synthesis. GenScript injected resultant protein into host animals, collected serum for testing, and affinity purified the product using additional target protein bound to a column. Antibody staining was performed as described previously for Drosophila and butterfly tissues^54^. Staged pupal wings and retinas were dissected and fixed 48 hours post-pupation. The Rabbit-anti-Bar antibody was used at 1:100, followed by secondary antibody staining with AlexFluor-555-anti-Rabbit secondaries at 1:500 and counterstaining with DAPI. Images were captured using standard confocal microscopy on a Leica SP5.

### CRISPR/Cas9 knockouts

The guide-RNA (gRNA) sequences were generated using the protocol described in Perry et al. 2016. Viable Cas9 target-sites were located by manually looking for PAM-sites (NGG) in the exon region of *BarH-1*. Uniqueness of the target regions was confirmed using a NCBI nucleotide blast (ver. 2.5.0+ using blastn-short flag and filtering for an e-value of 0.01) against the *C. crocea* reference genome. gRNA constructs were ordered from Integrative DNA Technologies (Coralville, Iowa, USA) as DNA (gBlocks). Full gRNA constructs had the following configuration: an M13F region, a spacer sequence, a T7-promotor sequence, the Target specific sequence, a Cas9 binding sequence, and finally a P505 sequence. Upon delivery, gBlocks were amplified using PCR to generate single-stranded guide RNA (sgRNA). For each gBlock, four 50ul reactions were conducted using the M13f and P505 primers and Platinum Taq (Invitrogen cat. 10966-034). The four reactions were then combined and purified in a Qiagen Minelute spin column (cat. 28004, Venlo, Netherlands). The resulting template was transcribed using the Lucigen AmpliScribe T7-flash Transcription Kit from Epicentre/Illumina (cat. ASF3507, Madison, WI, USA) followed by purification via ammonium acetate precipitation. Products were resuspended with Qiagen buffer EB, concentrations were quantified by Qubit and further diluted to 1000 ng/μl. They were then mixed with Cas9-NLS protein (PNA Bio, Newbury Park, CA, USA) and diluted to a final concentration of 125-250 ng/μl. *C. crocea* females (n > 40) from Aiguamolls de l’Empordà, Spain were captured and kept in morph-specific flight cages in the lab at Stockholm University where they oviposited on alfalfa *(Medicago sativa)*. Eggs were collected between 1-7h post-laying and sterilized in 7% benzalkonium chloride for ~5 minutes before injection. Injections were either at a concentration of 125 or 250 ng/ul and conducted using a M-152 Narishige micromanipulator (Narishige International Limited, London, UK) with a 50 ml glass needle syringe, with injection pressure applied by hand via a syringe fitting.

### CRISPR/Cas9 validation

To validate the mutation, Cas9 cut sites were PCR-amplified and a ~370bp region, centered on the intended cut site were sequenced using Illumina MiSeq 300bp paired-end sequencing. Primers were designed using Primer3 (http://biotools.umassmed.edu/bioapps/primer3_www.cgi). DNA was isolated from KO-individuals using KingFisher Cell and Tissue DNA Kit from ThermoFisher Scientific (N11997) and the robotic Kingfisher Duo Prime purification system. DNA quality and quantity were assessed via a Nanodrop 8000 spectrophotometer (Thermo Scientific, MA, USA) and a Qubit 2.0 Fluorometer (dsDNA BR; Invitrogen, Carlsbad, CA, USA). Aliquots were then taken and diluted to 1ng/ul before amplifying the region over the cleavage-site. Sequences were amplified and ligated with Illumina adapter and indexes in a two-step process following the protocol provided by Science for Life Laboratories (Stockholm, Sweden) and Illumina. First, we amplified the ~370bp long sequence around the cut sites and attach the first Illumina adapter, onto which we later attach Illumina handles and Index using a second round of PCR (Accustart II PCR Supermix from Quanta Bio [Beverly, MA, USA], settings 94C x 2 min followed by 40 cycles of 94 C x 30 sec + 60 C x 15 sec + 68 C x 1 min followed by 68 C x 5 min). PCR products were purified using Qiagen Qiaquick (Cat. 28104). Concentration and quality of the product were assessed via Nanodrop and gel electrophoresis. DNA was diluted to ~0.5ng/ul and then the unique double indices were attached by the second round of PCR (same protocol as above).

The final PCR products were purified again using Qiaquick spin columns and concentration and size was assessed using Qubit fluorometer and gel electrophoresis. All samples were then mixed at equal molarity and sent for sequencing at Science for Life Laboratories (Stockholm, Sweden). Sequences were aligned to their respective fragments (area surrounding cut site) using SNAP (ver. 1.0beta18)^55^, identical reads were clustered using the collapser utility in Fastx-Toolkit (http://hannonlab.cshl.edu/fastx_toolkit/). Sequences containing deletions were extracted and the most abundant sequences containing deletions were selected for confirmation of deletion in the expected region.

### Electron Microscopy

To quantify pigment granule differences between Alba and orange individuals pieces of the forewing were mounted on aluminum pin stubs (6mm length) with the dorsal side upwards. Samples were coated in gold for 80 seconds using an Agar sputter coater and imaged under 5 kV acceleration voltage, high vacuum, and ETD detection using a scanning electron microscope (Quanta Feg 650, FEI, Hillsboro, Oregon, USA). To quantify pigment granules within the photos we selected images from the same magnification and randomly placed three 4 μm^2^ squares on the images. We counted the number of pigment granules within each square and took the average, then conducted a two sample t-test in R. To quantify pigment granule differences between KO and wild type regions in our CRISPR KO mosaic individuals, a biopsy hole punch a 2mm in diameter circle was used to cut out one piece mostly containing white scales and one piece with mostly orange scales. These pieces were first photographed using a Leica EZ4HD stereo microscope in order to allow us to confirm the color of each scale once they were covered with gold sputter. Five white and five orange scales were then selected and the granules from a 4μm^2^ square were counted from each of those scales and a two sample t-test was then conducted in R.

### Lipid Analysis

Wild caught *C. crocea* Alba females (Catalonia, Spain) oviposited in the lab. Eggs were moved into individual rearing cups and split between two temperature treatments (hot: 27°C and 16 hour day length during larval and pupal development, cold: reared at 22 °C with a 16 hour day length during larval development and 15°C with a 16 hour day length during pupal development). Once pupated, individuals were checked a minimum of every 12 hours. Upon eclosion adults were stored at 4 °C until the next day to provide time for meconium excretion. Butterflies were not allowed to feed before dissection. Body weight was taken using a Sauter RE1614 scale before dissection. Total lipids were extracted using the Folch method^56^ according to the procedures outlined in Woronik et. al. 2018^11^. HPTLC was conducted as described in Woronik et. al. 2018^11^. In brief, 5 μl of the sample lipid extract was applied on a silica plate with a Camag Automatic TLC Sampler 4 (Camag, Muttenz, Switzerland). After the silica plate developed it was scanned with a Camag TLC plate scanner 3 at 254 nm using a deuterium lamp with a slit dimension of 6 × 0.45 mm and analyzed with the Win-CATS 1.1.3.0 software. Peaks representing the four major neutral lipid classes (diacylglycerols, triacylglycerols, cholesterol and cholesterol esters) were identified by comparing their retention times against known standards. Then the peak areas were integrated and the amount of lipid within each class was calculated using the formula: pmol_sample_ = (Area_sample_ / Area_standard_) x pmol_standard_. The total lipid content (nmol per abdomen) was calculated as a sum of pmol contents of all neutral lipid classes. For the statistical analyses this value was regressed against abdomen weight and standardized residuals (i.e. mass-corrected storage lipid amount) and were subsequently used as dependent variable.

### Transcriptome assembly and differential expression analysis

Offspring from a wild caught Alba female from Catalonia, Spain were reared at Stockholm University. When larvae reached the fifth instar they were checked at least every six hours and the pupation time of each individual was recorded. Tissue was collected between 82% and 92% of pupal development. Pupae were dissected in PBS solution, and the abdomen and wings were flash frozen in liquid nitrogen and stored at −80 °C. RNA was extracted from the abdomen and wing tissues using Trizol. RNA quality and quantity was assessed using a Nanodrop 8000 spectrophotometer (Thermo Scientific) and an Experion electrophoresis machine using the manufacturer protocol (Bio-Rad, Hercules, CA). Library preparation (Strand-specific TruSeq RNA libraries using poly-A selection) and sequencing (101 bp PE HiSeq2500 - high output mode) was performed at the Science for Life Laboratories (Stockholm, Sweden). In total 16 libraries were sequenced (4 orange and 4 Alba individuals - wings and abdomen from each individual). Raw data was cleaned and reads from all libraries were used in a de novo transcriptome assembly (Trinity version trinityrnaseq_r2013_08_14 with default parameters)^57^. To reduce the redundancy among contigs and produce a biologically valid transcript set, the tr2aacds pipeline from the EvidentialGene software package^58^ was run on the raw Trinity assembly. The sixteen RNA-Seq libraries were mapped to the resulting transcriptome using NextGenMap v0.4.10 (-i 0.09)^47^. SAMTOOLS v1.2^48^ was then used to filter (view -f 3 -q 20), sort and index the sixteen bam files. SAMTOOLS v1.2^48^ idxstats was then used to calculate the read counts per gene for each of the sorted bam files. These counts were then joined in a CSV file using an in-house pipeline and csvjoin^59^. A differential expression analysis was conducted in EdgeR^60^. A Benjamini Hochberg correction was applied to the raw p values to correct for false discovery rate and differentially expressed genes were called (adjusted p value <0.05). eggNOG-mapper (v.1)^61^ was used with default settings to functionally annotate the transcriptome. The R package topGo^62^ was used to conduct a gene set enrichment analysis on genes that exhibited > 1 or < −1 log fold change in the differential expression analysis.

